# Expanding the HIF-2α proteomic interactome with TurboID proximity labeling

**DOI:** 10.64898/2026.10.22.760981

**Authors:** Christine L. Faunce, Amina Khasanova, Ruofan Yu, Hua Huang, Francisca N. De Luna Vitorino, Michael B. Gilbert, Khoa A. Tran, Katherine A. Alexander, Benjamin A. Garcia, Charly R. Good, Shelley L. Berger

## Abstract

The transcription factor HIF-2α is a central regulator of cellular adaptation to hypoxia, orchestrating gene expression programs that control angiogenesis, erythropoiesis, and metabolic reprogramming, and is famously overexpressed in cancers such as clear cell Renal Cell Carcinoma (ccRCC). While the gene targets of HIF-2α have been extensively characterized, the composition and dynamics of its protein interaction network remain less well defined. In this study, we sought to systematically characterize proteins associated with HIF-2α using two distinct proteomic approaches, enabling the identification of both stable binding partners and transient or indirect interactors. We integrated affinity purification–mass spectrometry with proximity labeling strategies, thus expanding the existing interactome of HIF-2α to include uniquely identified proteins and complexes such as members of the nuclear pore complex. Across both immunoprecipitation mass spectrometry (IP-MS) and TurboID-MS, we identified 888 and 1,048 proteins, respectively. We applied stringent filtering criteria to refine both datatsets, yielding a dataset of 329 HIF-2α interacting proteins, approximately 75% of which have yet to be documented as HIF-2α interactors. We queried this set across publicly available cancer genomics and proteomics datasets and detected expression of our interactors in breast cancer and ccRCC tumor proteomes. These findings led us to identify GRHL2 and KIF4A as proteomic interactors that contribute to poor patient survival in breast cancer. Taken together, we herein provide a comprehensive resource of HIF-2α interacting proteins which provide insights into the mechanisms by which HIF-2α drives disease progression.

**Highlights:**

- 329 proteins were identified through immunoprecipitation and TurboID proximity labeling coupled with mass spectrometry based proteomics with 47 proteins commonly identified across both methods
- Interactome is largely composed of RNA binding and transcription factor binding facilitators
- Expression of a subset of HIF-2α interacting proteins are predictive of poor patient survival in breast cancer

## Introduction

HIF-2α, enocded by the gene *EPAS1*, is a transcription factor whose context-dependent activation regulates cellular adaption to hypoxia^1,2^. HIF-2α is one of three mammalian alpha subunits in the hypoxia inducible factor (HIF) family and shares structural and topological similarity with HIF-1α however, accumulating evidence suggests that it governs distinct transcriptional programs in a context-dependent manner^3,4^. The transcriptional program of HIF-2α regulates biological processes ranging from development^5^, angiogenesis^6^, erythropoesis^7^, metabolism^8^, and stem cell maintenance^2^. Dysregulated HIF-2α signaling is a major driver of tumor progression, most notably in clear cell Renal Cell Carcinoma (ccRCC), and its role in breast cancer disease progression is gaining attention^9–12^.

Protein–protein interactions are fundamental to transcription factor activity including mediating chromatin engagement, recruitment of coactivators, and signal integration. Appropriate function of HIF-2α requires heterodimerization with HIF-1β, encoded by the gene *ARNT*^13^, liscensing translocation into the nucleus where HIF-2α and HIF-1β can bind hypoxia repsonse elements and direct transcriptional output^14^. However, comprehensive investigation of the HIF-2α interactome is incomplete and current interactions have not been surveyed in breast cancer relevant samples, limiting our understanding of how HIF-2α coordinates disease progression.

Mass spectrometry–based proteomics offers a powerful approach to define such networks and has helped to elucidate HIF-2α’s interactome in melanoma^15^ and HeLa^16^ cell lines using epitope tag affinity-based pulldown methods. Affinity-based immunoprecipitation coupled to mass spectrometry (IP-MS) robustly identifies stable binding partners but cannot necessarily capture transient or weak interacting partners. In this study, we sought to utilize current proteomic approaches to expand the protein interaction landscape of HIF-2α using TurboID proximity dependent labeling. IP-MS and TurboID-based proximity labeling have complementary limtations. IP-MS can lose weak or transient interactions during stringent washing and fractionation. In contrast, TurboID captures these interactions in living cells but can only label proteins with accessible residues near the bait, so it may miss proximal proteins buried within sterically constrained complexes. Combining the two approaches therefore provides broader coverage of the HIF-2α interactome. This study establishes a systematically generated and well-annotated resource of HIF-2α–associated proteins, providing a foundation for future mechanistic and therapeutic investigations. Furthermore, we integrate our data with publicly available cancer genomic and survival data to begin understanding the biological signficiance of these interacting proteins.

## Experimental Procedures

### HIF-2α construct creation and stable cell line generation

A new polyclonal population of MCF7 cells expressing VHL degradation-resistant HIF-2α was generated following the previously published procedure from our laboratory.^17^. TurboID-fused HIF-2α constructs were generated using a Hi-Fi cloning strategy. The TurboID ligase was inserted on the N-terminus of the HIF-2α coding sequence and linked together by three repetitive SV40 nuclear localization signals (PKKKRKV). The TurboID sequence was PCR amplified using Hi-Fi strategy-based forward (5’-caaggacgacgatgacaagggatccatgaaagacaatactgtgcctc-3’) and reverse (5’-gatctgactgcggctcctcGGATCCCATcaccttcctcttcttctt-3’) primers and the Q5 High-Fidelity 2X Master Mix (NEB M0544S). The PCR product was run on a 1% agarose gel and purified using the Zymo Clean Gel DNA Recovery Kit (Zymo D4008). The HIF-2α overexpression backbone was linearized by restriction enzyme treatment (specifically, BAMHI-HF). 50ng of the backbone was used for the HiFi reaction giving a backbone:insert ratio of 20:1. The backbone and vector came together using the NEBuilder HiFi DNA Assembly Master Mix (NEB E5520S) in a 20uL reaction. The resulting product was transformed in stbl3 competent cells and whole plasmid sequencing using plasmidsaurus confirmed successful molecular cloning. All plasmids disclosed within this manuscript are available upon request.

Stable cell lines were created via lentivirus generation in low passage HEK293T cells (CRL-11268; ATCC) using pPAX2, pVSV-G, and lipofectamine2000 according to published protocols^18,19^. MCF7 cells were infected with a 1:2 ratio of virus containing media and MCF7 growth media. To induce HIF-2α-TurboID expression, doxycyclin (Sigma, D3447-500MG) was dissolved in water to a 30mM stock solution and added to MCF7 cells at a final concentration of 3uM. Cells tested negative for mycoplasma contamination.

### Immunoprecipitation-Mass Spectrometry Sample Preparation

HIF-2α-expressing MCF7 cells were grown in 15cm^2^ plates in DMEM with 10% FBS and 1% antibiotics until 90% confluent. The night before collection, bead-antibody conjugates were assembled for overnight incubation at 4C. In low-adhesion microcentrifuge tubes, a 1:1 mixture of protein A (ThermoFischer Scientific 10002D) and protein G dynabeads™ (ThermoFischer Scientific 10004D) for a total volume of 30μL per sample were mixed. The beads were prewashed and then 5μg of HIF-2α antibody (Rabbit Novus NB100-122) or 5μg of IgG (abcam 6715 lot GR3382828-11) antibody was added per sample in 1mL of 1% BSA in dPBS.

Cells were treated with doxycycline for 6 hours and then washed with ice-cold dPBS twice. Cells were resuspended in 5mL hypotonic buffer (10 mM Tris-HCl, pH 8.0, 5 mM KCl, 1.5 mM MgCl, 1 mM DTT, phosphatase inhibitors, and EDTA-free protease inhibitors)^20^ to facilitate nuclear isolation. Cells were incubated on ice for 15 minutes and subsequently scraped with a cell lifter to rupture the cell membrane and then pelleted by centrifugation at 2000RPM for 5 minutes at 4C. The pellet, containing the nuclei, was gently washed once with hypotonic buffer and then incubated with 500uL of RIPA lysis buffer supplemented with 1X HALT protease and phosphatase inhibitor (Invitrogen NP0001-02). The samples were incubated on ice for 1 hour, vortexing every 10 minutes. The samples were subsequently digested with benzonase at a 1:500 dilution (Millipore 70746-3) and magnesium chloride (stock: 100mM, working: 1mM) and incubated at room temperature for 10 minutes. Samples were centrifuged at 16,000RPM for 10 minutes at 4C and the supernatant was collected as the nuclear fraction.

Antibody-bead conjugates were washed in a gradient of blocking solution (1% BSA in PBS) to IGEPAL over 4 washes rotating at 4C for 5 minutes: 100% blocking solution, 50% blocking solution and 50% IGEPAL lysis buffer twice, 100% IGEPAL lysis buffer. After the final wash, beads were resuspended in 100uL IGEPAL lysis buffer per sample and redistributed to individual low-affinity microcentrifuge tubes. Nuclear lysates of 1mg/mL concentration in 500uL of IGEPAL buffer were added to the antibody-bead conjugates (600uL total volume) and incubated overnight at 4C while rotating. The next morning, the bead-lysate conjugates were bound to a magnetic stand for the collection of the flow-through. The bead-antibody-lysate conjugates were washed with 500uL of IGEPAL lysis buffer twice while rotating at 4C. This was followed by two washes with 500uL of PBS while rotating at 4C. After the final wash, the bead-antibody-lysate conjugates were resuspended in 100uL of 50mM ammonium bicarbonate (stock: 100mM). Disulfide bonds were reduced with 5.5uL of freshly prepared 100mM DTT for a final concentration of 10mM and samples were incubated at 55C while shaking at 1200RPM for 20 minutes. Samples were alkylated with 25mM iodoacetamide (stock: 250mM) and incubated in the dark for 45 minutes. Samples were digested with 0.8uL of sequencing-grade trypsin (Promega V5113).

### TurboID-Mass Spectrometry Sample Preparation

HIF-2α-TurboID-expressing MCF7 cells were grown in 15cm^2^ plates in DMEM with 10% FBS and 1% antibiotics until 90% confluent. Cells were treated with doxycycline for 6 hours and immediately refreshed with media containing biotin (Sigma B4501-1G; stock 100mM; working 50uM) for 15 minutes. Cells were collected, lysed, and enriched according to previously described conditions^21^. Cells werewashed with ice-cold dPBS five times. After the last wash, cells were scraped and pelleted by centrifugation at 300RPM for 3 minutes at 4C. The pellets were lysed with 500uL RIPA lysis buffer (ThermoFischer 89900) supplemented with 1X HALT protease and phosphatase inhibitor and incubated on ice for 10 minutes. The cell lysates were clarified by centrifigation at 13,000RPM for 10 minutes at 4C. 83uL of pre-washed streptavidin beads (Thermo Pierce 88817) were added to 1mg of protein lysate in 500uL of RIPA lysis buffer and the resulting bead-lysate slurry was incubated rotating overnight at 4C. The following morning, the bead-lysate conjugates were bound to a magnetic stand for the collection of the flow-through. The bead-lysate conjugates were washed with 500uL of 50mM Tris-HCl (pH 7.5) twice while gently rotating at 4C. This was followed by two washes with 500uL of 2M urea in 50mM Tris-HCl (pH 7.5). The bead-lysate conjugates were incubated with 80 μL of 2M urea in 50 mM Tris-HCl (pH 7.5) containing 1 mM DTT and 0.4 μg trypsin (Promega V5113) at 25C for 1 hour while shaking at 1,000RPM. The supernatant was collected from this digestion and combined with two subsequent washes with 60μL of 2 M urea in 50 mM Tris HCl (pH 7.5) for a total supernatant volume of 200uL. Freshly prepared DTT was added to a final concentration of 4mM and incubated at 25C for 30 minutes while shaking at 1,000RPM to reduce the disulfide bonds. The samples were incubated with 10mM iodoacetaminde at 25C while shaking for 45 minutes. An additional 0.5ug of trypsin was added to the sample for overnight digestion at 25C while shaking. The next morning, samples were acidified to a pH less than 3 with the addition of FA.The resulting peptides were dried using a SpeedVac centrifuge. The peptides were desalted with the self-packed C18 stage tip. The purified peptides were then dried and reconstituted in 0.1% formic acid.

### C18 Stage Tip Desalting

Following proteolytic digestion, all peptide samples were desalted and concentrated using C18 reversed-phase StageTips based on previously described StageTip protocols^22^. C18 StageTips were conditioned with 100 μL of LC-MS-grade methanol and centrifuged at 2,000RPM for 1 min at room temperature. The StageTips were subsequently equilibrated twice with 50 μL of 0.1% trifluoroacetic acid (TFA), with centrifugation at 2,000RPM for 1 minute following each addition. Samples were resuspended in 50 μL of 0.1% TFA and loaded onto the equilibrated C18 StageTips by centrifugation at 2,000RPM for 1 minute. The StageTips were washed twice with 50 μL of 0.1% TFA, centrifuging at 2,000RPM for 1 minute after each wash, to remove salts and other hydrophilic contaminants. Peptides were subsequently eluted into clean microcentrifuge tubes using two sequential 50μL aliquots of 60% acetonitrile (ACN), with centrifugation at 2,000RPM for 1 min following each elution. The combined eluate was concentrated to dryness using a SpeedVac concentrator at medium speed for approximately 30 min. Dried peptides were resuspended in 20μL of 0.1% formic acid (FA) prior to LC-MS/MS analysis.

### LC-MS/MS and Data Analysis

An LC-MS/MS system consisting of a Vanquish Neo UHPLC coupled to an Orbitrap Exploris 240 or Ascend (Thermo Scientific) was used for peptide analysis. Separation of peptides was carried out on an Easy-Spray™ PepMap™ Neo nano-column (2 µm, C18, 75 µm X 150 mm) at room temperature with a mobile phase. The chromatography conditions consisted of a linear gradient from 2 to 32% solvent B (0.1% FA in 100% ACN) in solvent A (0.1% FA in water) over 75 minutes and then 45 to 98% solvent B over 15 minutes at a flow rate of 300 nL/min. The mass spectrometer was programmed for data-dependent acquisition (DDA) mode. The MS1 was acquired over a mass range of 295-1100 m/z with a resolution of 120,000, an automatic gain control (AGC) target of 300%, and a maximum injection time of 50 ms.

For the MS2, the top 20 most intense ions were selected for MS/MS by high-energy collision dissociation (HCD) at 30 nominal collision energy (NCE), with a resolution of 15,000, AGC target of 75%, and maximum injection time of 60 ms. Raw files were processed using FragPipe^23^. Carbamidomethyl (C) was set as a static modification while oxidation (M) and acetylation (protein N-terminus) were selected as variable modifications.

The resulting list of unique peptides were assigned to proteins, and this list was further filtered to exclude proteins mapped to only one unique peptide, common contaminant proteins as well as non-nuclear proteins. Contaminant filtering was performed against the CC156 database^24^ which includes data from a HEK293 cell line expressing GFP (Green Fluorescent Protein) targeted to the nucleus. Non-nuclear protein filtering was performed using the Human Protein Atlas’ subcellular localization annotations^25^. IP-MS data were filtered to only include proteins which were identified across at least two of four biological replicates and had a logFC value greater than 1.5 over IgG control.

This data is currently being uploaded to publicly available data repository.

### Immunofluorescence and Analysis

Cells were washed with 4% paraformaldehyde diluted in nuclease-free (NF)-PBS solution and incubated for 10 minutes. Cells were then permeabilized by 0.2% Triton X-100 (ThermoFischer A16046.AE) diluted in NF-PBS for an additional 10 minutes at room temperature. Cells were stained in primary antibody (1:100) overnight at 4C. Cells were washed three times with NF-PBS and stained in secondary antibody (1:1000) for 1 hour at room temperature protected from light. Cells were washed three times with PBS and then mounted. Cells were imaged on a Nikon Eclipse Ti2 using Nikon NIS software.

The following antibodies and fluorescent compounds were used in this study: Invitrogen D1306 (DAPI) was used to stain double-stranded DNA to label the nucleus; Novus Bio NB100-122 for primary staining of HIF-2α; Jackson Immunoresearch 016-600-084 Alexa Fluor® 647 Streptavidin; ThermoFischer A-11008 for anti-Rabbit secondary antibody staining.

The percentage of HIF-2α-positive cells was quantified using CellProfiler. Colocalization between the DAPI and HIF-2α channels was measured for each field of view. For each biological replicate, the mean percentage of HIF-2α-positive cells was calculated across at least 10 fields of view and treated as a single technical replicate. At least three biological replicates, defined as independent imaging wells, were analyzed. Fluorescence line-profile analyses were performed using FIJI/ImageJ software.

### IP-Western Immunoblots

During the both pulldown protocols, input, flow-through, and IP/bead samples were collected for IP-Western. Samples were prepared in a 6.25:5:2 ratio of sample:sample buffer:reducing reagent and denatured at 95C for 10 minutes. Samples were loaded into NuPAGE™ 4-12% Bis-Tris Mini Protein Gels(ThermoFischer NP0336) with 1X MOPS buffer (ThermoFischer NP000102). The membrane was transferred into 0.45um nitrocelluose membrane (Invitrogen LC2001) and blocked with 5% bovine serum albumin (BSA). Primary antibody was incubated overnight at 4C. The next morning, the membranes were washed three times with 1X TBST and then incubated with secondary HRP-conjugated antibodies for 1 hour at room temperature. Membranes were washed three times with 1X TBST at 10 minute intervals. HRP signal was activated using the SuperSignal™ West Pico PLUS Chemiluminescent Substrate (ThermoFischer 34580).

The following antibodies and HRP-conjugated compounds were used in this study: Novus Bio NB100-122 for primary staining of HIF-2α; Cell Signaling D16H11 for primary staining of GAPDH; Santa Cruz sc-55526 for primary staining of ARNT/HIF-1b; Biorad 1706515 for secondary staining of anti-Rabbit primary antibodies.; Biorad 1706516 for secondary staining of anti-Mouse primary antibodies.

### cBioPortal Database Analysis

Genomic alteration and clinical survival analyses were performed using the cBioPortal for Cancer Genomics. Data was queried from the Breast Invasive Carcinoma (TCGA, PanCancer Atlas and METABRIC, Nature 2012 and Nat Commun 2016) datasets, encompassing a total of 1,084 and 2,509 patient samples, respectively. To evaluate the co-occurrence of copy-number amplifications, an OncoPrint analysis of the TCGA – PanCancer Atlascohort (n = 1,084) was performed. The query was restricted to amplification events using Onco Query Language (OQL) syntax (GENE: AMP). Genes-encoding our interacting proteins were altered in 632 patients.

Gene expression and clinical data were obtained from the same datasets. For each interactor identified in our study, a univariable Cox proportional hazards model was fitted in each cohort separately, relating overall survival to standardized expression, and the Wald z statistic was recorded. Expression of each gene was standardized as a within-cohort z-score. Patients with missing expression or survival data, or with a follow-up time of zero, were excluded. Within each cohort, patients were classified as high or low for EPAS1 and for the partner gene (GRHL2 or KIF4A) according to whether expression was above the cohort median, giving four groups (both low, EPAS1 high only, partner high only, both high). No optimized cut points were used. The endpoint was overall survival in both cohorts. Survival curves were estimated by the Kaplan–Meier method and compared across the four groups using a log-rank test. The both-high and both-low groups were also compared directly using a log-rank test, with the hazard ratio and 95% confidence interval estimated from a Cox proportional hazards model. To test whether the joint effect exceeded the sum of the individual effects, a Cox model containing EPAS1, the partner gene, and their product term was fitted using continuous z-scores, and the Wald P value for the interaction term is reported. For display, curves were truncated at 250 months (METABRIC) or 150 months (TCGA); all statistics were computed on the full follow-up. Analyses were performed in R using the survival and tidyverse packages.

### CPTAC Database Analysis

HIF-2α interactors were defined as any protein identified across either method after peptide, nuclear, and CRAPome filtering. As a background set, 2,000 protein-coding genes were randomly sampled without replacement, excluding the interactors, and using a fixed random seed (1,000). The same background set was used in all analyses. Protein abundance data was taken from three breast cancer references and one ccRCC reference. For each patient, the detection rate of each gene set was calculated as the percentage of genes in the set whose protein product was detected, separately for tumor and matched normal tissue. The paired difference (tumor minus normal) was calculated per patient and is reported in percentage points. Dashed lines indicate the median paired difference across patients. For each protein quantified in both tumor and matched normal tissue in 10 pairs, the log2 fold change (tumor/normal) was calculated within each pair and summarized across patients as the median. This yielded 321 interactors and 1,053 background proteins. Distributions were compared using a two-sided Wilcoxon rank-sum test. Box plots show the median and interquartile range, with whiskers extending to 1.5 × the interquartile range. The percentage of proteins "up in tumor" refers to those with a summarized log2 fold change above zero.

## Results

### Experimental workflow and validation of orthogonal proteomic approaches for mapping the HIF-2α interactome

HIF-2α is a transcription factor, indicating the potential for numerous interactions, both direct and indirect (or transient). To capture interacting proteins of various interaction distances, we performed an affinity-based pulldown of HIF-2α alongside proximity labeling of HIF-2α fused with TurboID. Immunoprecipitation pulldown of HIF-2α is expected to measure direct interactors, whereas TurboID conjugated HIF-2α is expected to label both direct (as validation of IP-MS) and indirect/proximal interactors (**Figure 1A**). To facilitate the examination of HIF-2α interacting protein partners, we generated HIF-2α expressing and HIF-2α-TurboID expressing constructs, and stably expressed them in MCF7 breast cancer cells, a cell line that expresses low levels of endogenous HIF-2α in normoxic conditions due to proteasomal degradation triggered by von Hippel Lindau (VHL) mediated ubiquitination^9,10,26^. Importantly, the HIF-2α expression constructs were engineered to contain mutations at amino acid residues 405 and 531 which prevents HIF-2α recognition and degradation by VHL, allowing stable protein expression^17,27^. Similar cell lines expressing stabilized mutants of HIF-2α in MCF7 cells have been used to uncover how HIF-2α regulates nuclear speckle association and distinct transcriptional programs in breast cancer cells^17^. This led us to question how HIF-2α coordinates with other proteomic partners to drive tumorigenesis in breast cancer.

**Figure 1:**
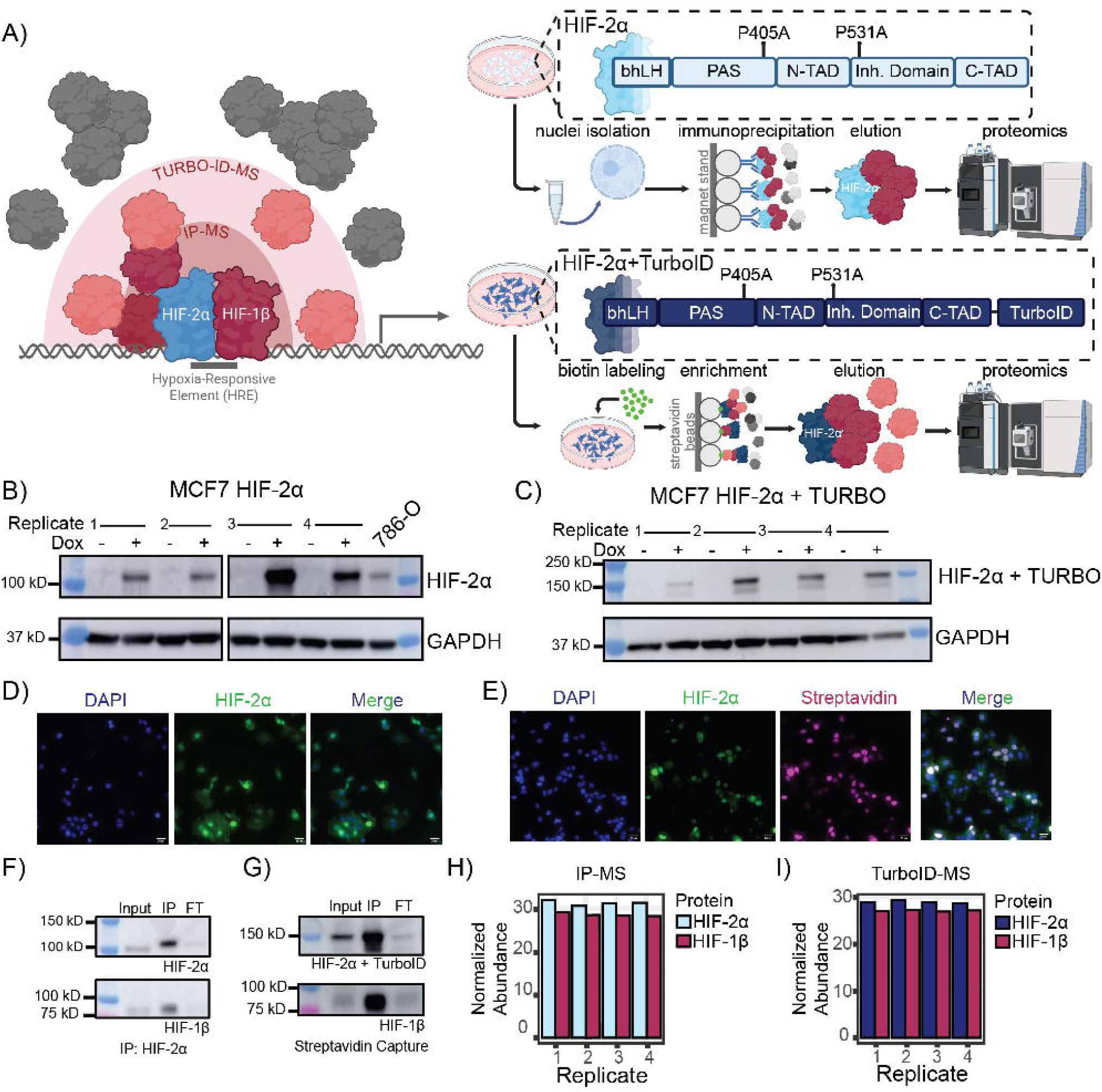
Experimental workflow and validation of orthogonal proteomic approaches for mapping the HIF-2α interactome. **A)** Cartoon schematic of both proteomic workflows: IP-MS used to label direct interactors and TurboID-MS used to label both direct and indirect interactors of HIF-2α. **B)** Western Blot image showing doxycycline-inducible expression of HIF-2α. 786-O cell lysate was run in the last lane as a molecular weight reference for HIF-2α. **C)** Western Blot of MCF7-HIF-2α-TurboID cell lysate with and without doxycycline treatment. HIF-2α runs at a higher molecular weight than in B) due to the conjugation to the TurboID sequence (∼35kDa) to HIF-2α. **D)** Immunofluorescence imaging of MCF7 HIF-2α cells after treatment with doxycycline. Primary antibody staining against HIF-2α shown in green. DAPI staining shown in blue. **E)** Immunofluorescence imaging of MCF7 HIF-2α-TurboID cells after treatment with doxycycline. Primary antibody staining against HIF-2α shown in green. DAPI staining shown in blue. Streptavidin-647 fluorophore shown in magenta. **F)** Western blot of HIF-2α immunoprecipitates from MCF7-HIF-2α cells, probed for HIF-2α and its dimerization partner HIF-1β. Remaining replicates in Figure S1G. **G)** Western blot of HIF-2α immunoprecipitates from MCF7-HIF-2α-TurboID cells, probed for HIF-2α and its dimerization partner HIF-1β. Remaining replicates in Figure S1H. **H)** Normalized abundance of HIF-2α and HIF-1β pulldown in IP-MS across four replicates. **I)** Normalized abundance of HIF-2α and HIF-1β pulldown in TurobID-MS across four biological replicates.

We first sought to validate our cell culture systems for proper expression and subcellular localization of HIF-2α. Western blot imaging of both cell lysates confirmed induction of HIF-2α signal upon treatment with doxycycline compared to cells without doxycycline treatment (**Figures 1B-C, S1A**). Immunofluorescence confirmed strong, nuclear expression of HIF-2α following doxycycline treatment in HIF-2α (**Figure 1D**) and HIF-2α-TurboID expressing cells (**Figure 1E**). The majority of the cells (>75% on average in each field of view) had a HIF-2α positive signal in the nucleus as measured by the colocalization of an anti-HIF-2α antibody and DAPI channel signal (**Figures S1B-C**). As an orthogonal validation of nuclear spatial distribution, we measured the fluorescence line profiles across single nuclei of MCF7 HIF-2α (**Figure S1D**) cell lines in the presence of doxycycline. A linear region of interest was drawn across a representative cell, spanning the cytoplasm and the nucleus which showed a sharp increase in HIF-2α fluorescence signal (green channel) at the nuclear boundary as marked by DAPI (blue channel). A similar analysis was performed to quantify the spatial distribution of biotinylated proteins upon biotin treatment to HIF-2α-TurboID cells (**Figure S1E**). As shown in the line profile plot, the fluorescence intensity of biotinylation signal via a Streptavidin-Alexa Fluor™ 647 Conjugate (dark red channel) sharply increased at the nuclear boundary and mirrored the intensity trajectory of both DAPI (blue channel) and HIF-2α (green channel). To assess optimal time for biotin labeling, we performed a time course of biotin treatment on doxycycline-induced HIF-2α-TurboID cells and used a streptavidin conjugated fluorophore to assess biotinylation over time^28^ (**Figure S1F**). A biotin labeling time of 15 minutes was used for subsequent experiments. These results confirm that viral transduction of cells carrying either HIF-2α or HIF-2α-TurboID was successful and led to the appropriate subcellular localization of protein expression.

To evaluate interaction fidelity for each replicate in both techniques, HIF-2α pulldown was compared to co-association of its established heterodimer partner, ARNT/HIF-1β^14^. In each approach, using Western Blot analysis, HIF-1β appeared proportionally enriched compared to HIF-2α levels (**Figures 1F-G** examples and **Figures S1G-H** all replicates), showing detection of direct interaction with a known associating factor required for transcriptional activity of HIF-2α. These samples were next analyzed by mass spectrometry, and we first evaluated the pulldown abundance of HIF-2α across all four biological replicates for each method. IP-MS yielded robust and reproducible measurements of HIF-2α protein abundance, reflecting the relative amount of HIF-2α detected in each sample, with normalized abundance values ranging from 30.7 to 32.1 (**Figure 1H**). TurboID-MS showed a comparable pulldown abundance across replicates, with pulldown abundances ranging from 28.6 to 29.3 (**Figure 1I**). Further, quantification of HIF-1β enrichment also showed consistency between replicates for both approaches (**Figures 1H-I**). Together, these results assure reliability of both approaches and permit further investigation of the results of new putative interacting partners of HIF-2α.

### Integrated IP-MS and TurboID-MS analyses identify a high-confidence core HIF-2α interactome

Confident in our methods to detect interactors of HIF-2α across both methods, we proceeded with systematic classification of the HIF-2α-associated proteome. We used five filtering steps to acquire a stringent dataset from our full proteomics data. As a first step, we removed any proteins that were mapped to a single unique peptide (**Figures S2A-B**). Density plots of log2-transformed abundances showed variability in intensity distributions across replicates; we therefore applied variance-stabilizing normalization (VSN) to all data, which tightened the spread of abundance values in both the IP-MS and TurboID-MS datasets (**Figures S2C-D**). While IP-MS samples underwent nuclei isolation, TurboID samples did not. To limit downstream analyses to nuclear-specific functions of HIF-2α, we next filtered against the Human Protein Atlas’ annotated list of nuclear proteins^25^. Of 888 proteins identified by IP-MS and 1,048 proteins identified by TurboID-MS, 23.9% and 31.5% were annotated as nuclear, respectively (**Figure S2E**). While filtering against the Human Protein Atlas is a strict and limiting filter, we were surprised by attrition of so many proteins. To confirm that this analysis performed as expected by removing any non-nuclear proteins, we performed Gene Ontology (GO) analysis on the excluded proteins. In both approaches, the removed proteins were most strongly enriched for translation and other cytoplasmic processes (**Figures S2F-G**). To eliminate non-specific background contaminants, proteins identified in both methods were filtered against the CRAPome database experiment CC156, which serves as a comparable negative control for nuclear samples^24^ (**Figure S2H**). We kept proteins that were identified in at least 2 biological replicates in both methods. Finally for the IP-MS, we filtered for proteins that were enriched over IgG controls, resulting in a dataset of 93 proteins, which included HIF-2α and HIF-1β (**Figure S3A**). These filtering steps resulted in final datasets of the following sizes: 93 IP-MS proteins and 283 TurboID-MS proteins.

Since interactomes of HIF-2α have been previously published in a melanoma cell line^15^ as well as the HeLa cell line^16^, we compared the proteins identified in their studies as well as across both of our methods. We found 6 of the 66 proteins from Steunou et al. overlapped between both of our datasets, including HIF-2α and its obligate binding partner, HIF-1β (**Figure S3B**). The remaining four overlapping proteins between Steunou et al. and our interactome include EMD, MED1, ARNT2, and MED12. An additional 9 proteins were identified in Steunou et al. and either our IP-MS or TurboID-MS datasets. Daly et al. evaluated normoxic and hypoxic binding partner differences of HIF-2α, and our dataset identified 76 of the same proteins (**Figure S3C**). Daly et al. classified their hits into three categories (hypoxic, normoxic, and oxygen independent), and we identified roughly the same proportion of binding partners across all three categories (4.3%, 5%, and 6% respectively) despite the lack of hypoxic treatment conditions in our experiments.

Next, to gain a deeper understanding of our HIF-2α interactors in breast cancer, we compared the intersection of our IP-MS and TurboID-MS datasets, yielding a total of 47 proteins that were detected in both approaches (**Figure 2A**; Fisher’s exact test, p < 0.0001). In contrast, 46 proteins were uniquely identified by IP-MS, while TurboID-MS captured a substantially larger set of 236 unique proteins. The increased coverage observed in the TurboID-MS dataset is consistent with its mechanism of labeling proteins in the spatial vicinity of the bait, including transient or weakly associated proteins that may not be retained under the more stringent conditions required for affinity purification^21,29^. Together, this intersection defines a core set of high-confidence interactors detected by both approaches, alongside distinct method-specific populations that may reflect differences in biochemical capture space.

**Figure 2:**
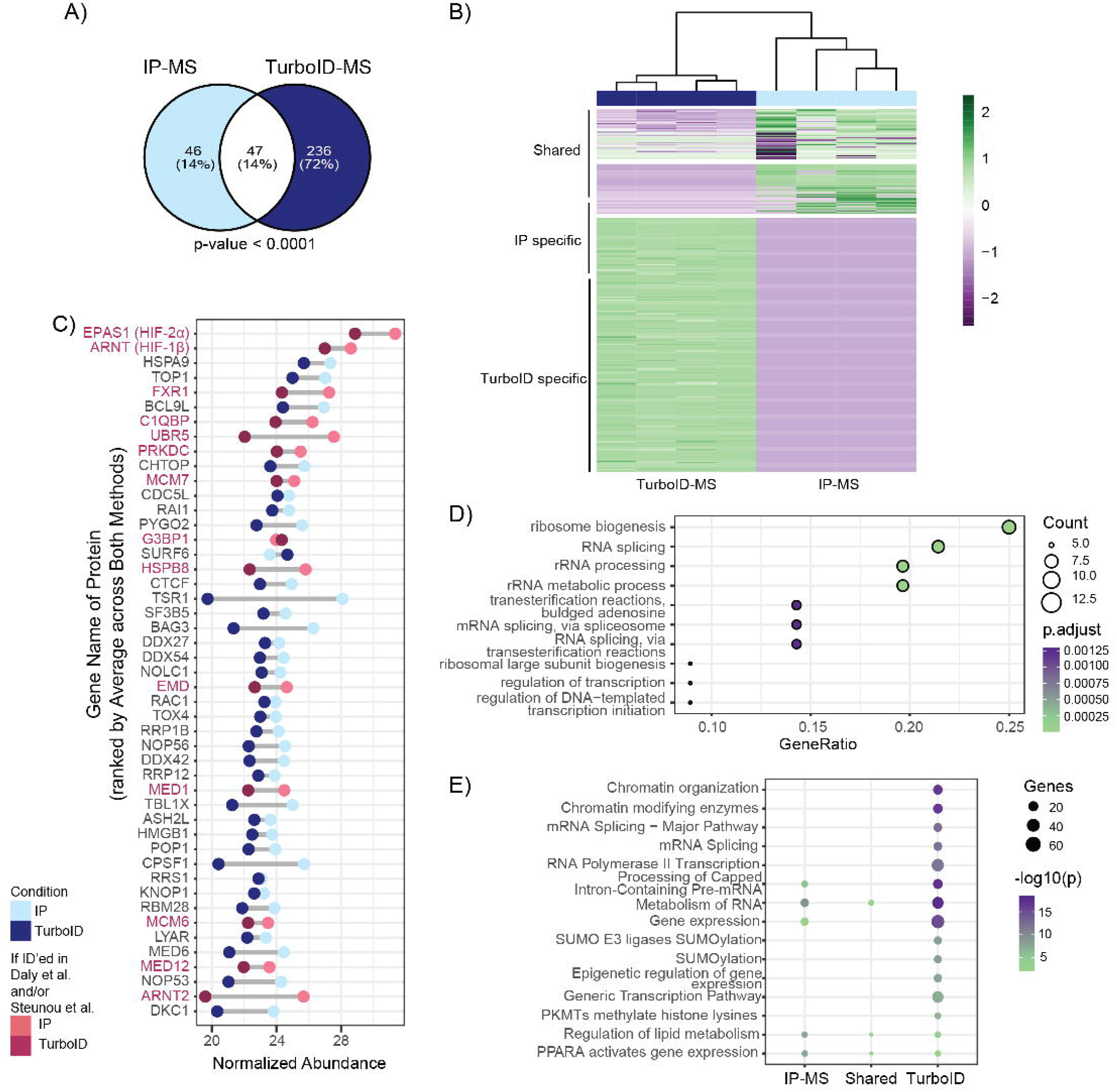
Integrated IP-MS and TurboID-MS analyses identify a high-confidence core HIF-2α interactome. **A)** Venn diagram illustrating the overlap between proteins identified by IP-MS and TurboID-MS. Statistical significance of the overlap was determined using a hypergeometric test. **B)** Hierarchical clustering heatmap of the all proteins identified across all IP-MS and TurboID-MS biological replicates. Protein abundances were VSN-transformed and row-scaled (Z-score) prior to clustering. **C)** Ranked abundance plots of proteins identified by both proteomic approaches. Proteins are ordered according to their average normalized abundance across IP-MS and TurboID-MS datasets, with individual measurements shown for each method. Proteins colored in red were found in Daly et al. and/or Steunou et al. **D)** Gene Ontology - Biological Process for the 74 shared proteins. **E)** Reactome pathway enrichment analysis of unique proteins identified by IP-MS, TurboID-MS, or both. Dot size represents the number of proteins assigned to each pathway, and color indicates enrichment significance (-log P value).

To further resolve these relationships, proteins were stratified into three classes: IP-MS– only, TurboID–only, and shared interactors. Hierarchical clustering of normalized abundance profiles across replicates with row-by-row z-score standardization provided a granular view of pulldown differences across all replicates (**Figure 2B**). Shared proteins exhibited relatively consistent detection patterns across both datasets, whereas IP-MS–only proteins showed enrichment primarily in immunoprecipitation-based capture conditions. In contrast, TurboID–only proteins formed a distinct cluster characterized by broader detection in proximity-labeling experiments, consistent with their assignment to the local proteomic environment of the HIF-2α bait rather than association in a potential stable complex. Together, these analyses highlight the utility of TurboID in expanding coverage to include spatially proximal components of the bait-associated proteome. It was unexpected that all proteins identified from IP-MS were not captured in TurboID-MS, but this could be an artifact of occluded biotinylation sites on interacting proteins or steric hinderance from the HIF-2α-TurboID fusion construct^30^.

Of the shared proteins, HIF-2α and its obligate binding partner, HIF-1β, were most abundantly pulled down across both detection methods, as expected (**Figure 2C**). The normalized abundance of each protein mostly agreed across detection methods, although the detection was often greater in IP-MS samples. However, since these sets of samples were not run at the same time, we do not glean biological significance from this finding. Of our 47 shared proteins, 14 were also identified in at least one of the two previously published HIF-2α interactomes^15,16^. StringDB analysis^31^ broadly identified RNA binding, and regulation of transcription as enriched functional categories from this subset of commonly identified interactors (**Figure S3D**). These shared proteins are enriched in Gene Ontology (GO) Molecular Function pathways related to nucleic acid binding and isomerase activity (**Figure S3E**) and GO Biological Process pathways related to ribonucleoprotein complex biogenesis and splicing (**Figure 2D**). Reactome analysis^32^ of the interactors found in IP-MS-only, TurboID-MS-only, and Shared revealed some overlapping and some distinct enrichment signatures for TurboID-identified proteins (**Figure 2E**). Across all three classes, mRNA processing and lipid metabolism categories were highly enriched. TurboID-only interactors uniquely enriched in chromatin organization, splicing-related SUMOylation, and epigenetic regulation of gene expression. The results from the Reactome analysis support the premise that a core set of interactors have been co-verified by pulldown across both methods while TurboID-MS expands the putative interactome.

### IP-MS identifies a HIF-2α interaction network enriched for transcriptional regulators

The distinct pathway enrichment profiles identified by Reactome analysis indicate that IP-MS and TurboID-MS capture different subsets of HIF-2α-associated proteins. We therefore performed separate downstream analyses of each interactome to better characterize the unique biological processes represented by each approach. We ranked all IP-MS detected proteins by average normalized abundance, which revealed a continuous distribution with HIF-2α and HIF-1β as two of the most abundant proteins recovered (**Figure 3A**), consistent with efficient bait enrichment. The top 30 highest-abundance interactors, ranked by average normalized abundance, were recovered with high reproducibility across replicates (**Figure 3B**). Notably, several of the highest-confidence interactors are RNA-binding proteins or components of RNA processing machinery, including the cleavage/polyadenylation factor CPSF1, the RNA-binding proteins FXR1 and CHTOP, and the ribosome biogenesis factor TSR1. The interactome also included known transcriptional co-regulators, such as the coactivator CARM1^33^. Notably, seven Mediator complex members were robustly pulled down with HIF-2α (**Figure 3C**). Mediator proteins were also captured by our TurboID pulldown (**Figure S3F**) as well as in the published melanoma cell line interactome publication^15^. To gain a broader, unbiased understanding of the IP-MS interactome, functional enrichment analysis of the significantly enriched proteins was performed through Gene Ontology Biological Process analysis (**Figure 3D**). This revealed an overrepresentation of biological processes ribonucleoprotein complex biogenesis, RNA processing, and transcription initiation and elongation. These findings suggest that, in addition to its established transcriptional machinery, HIF-2α may be associated with proteins involved in RNA metabolism and ribonucleoprotein assembly.

**Figure 3:**
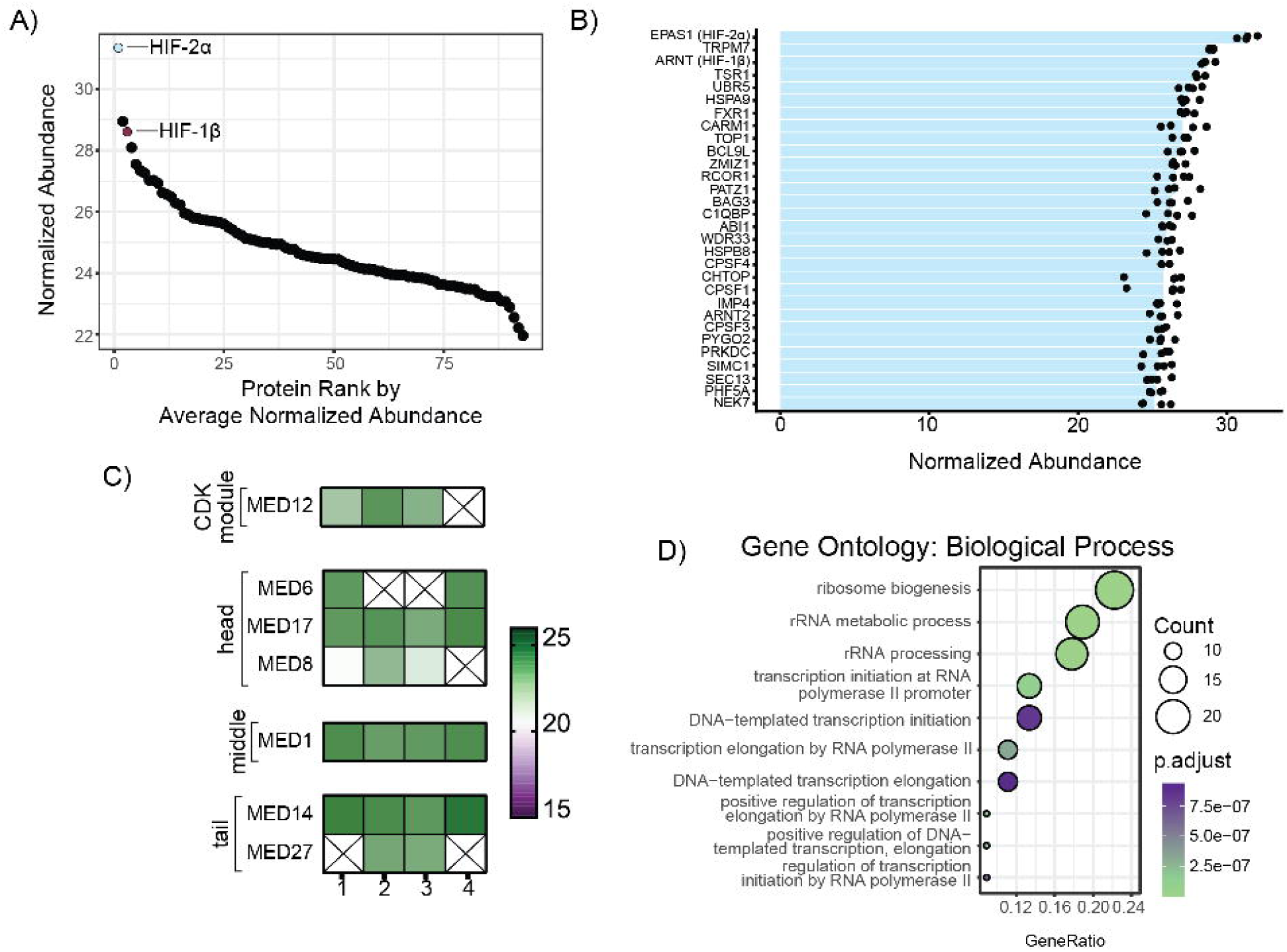
IP-MS identifies a HIF-2α interaction network enriched for transcriptional regulators. **A)** Rank plot displaying the abundance distribution of proteins identified in the HIF-2α IP-MS dataset. Proteins ranked by average normalized abundance. **B)** Top 30 ranked proteins identified by HIF-2α IP-MS are shown based on normalized abundance values. **C)** Heatmap depicting components of the Mediator complex groups by domain (head, middle, tail, and kinase-associated modules). **D)** Gene Ontology (GO) biological process enrichment analysis of IP-MS-identified HIF-2α-associated proteins. Dot size indicates the number of proteins contributing to each enriched term, while color represents adjusted p-value.

### TurboID-MS expands the HIF-2α interaction network to include mRNA export factors

Compared to IP-MS, TurboID-MS identified a substantially larger number of proteins, consistent with the ability of proximity labelling to capture both stable and transient protein associations. As observed in the IP-MS dataset, HIF-2α and its obligatory binding partner HIF-1β were among the most abundant proteins identified among 283 proteins (**Figure 4A**), confirming efficient biotinylation of proteins in the immediate vicinity of HIF-2α. The most abundant proteins recovered were HSP proteins^34^ as well as proteins involved in chromatin regulation and RNA metabolism, such as CHD4^11^ and TOP1^35^, which are reported regulators of HIF stability (**Figure 4B**). A prominent feature of the TurboID-MS dataset was the enrichment of nuclear pore complex (NPC) proteins. Multiple core scaffold nucleoporins, including NUP62, NUP93, NUP98, NUP153, and POM121C, together with peripheral nucleoporins (NUP88, NUP214, and NUP358/RANBP2) and nuclear transport-associated proteins (NXF1, RAE1, and SUMO1) were consistently detected at high abundance (**Figure 4C**). These proteins were not detected in our IP-MS pulldown (**Figure S3G**), suggesting that HIF-2α may transiently interact with components of the nuclear transport machinery, consistent with the continuous nucleocytoplasmic trafficking required for HIF signaling^36^. GO Biological Process analysis highlighted pathways related to RNA splicing, ribonucleoprotein complex biogenesis, mRNA processing, spliceosomal assembly, and ribosome biogenesis (**Figure 4D**). Collectively, these data indicate that the HIF-2α proximal proteome is enriched for proteins involved in transcriptional regulation, RNA processing, and nuclear transport, extending the interaction landscape identified by IP-MS and providing a broader view of the molecular environment surrounding HIF-2α.

**Figure 4:**
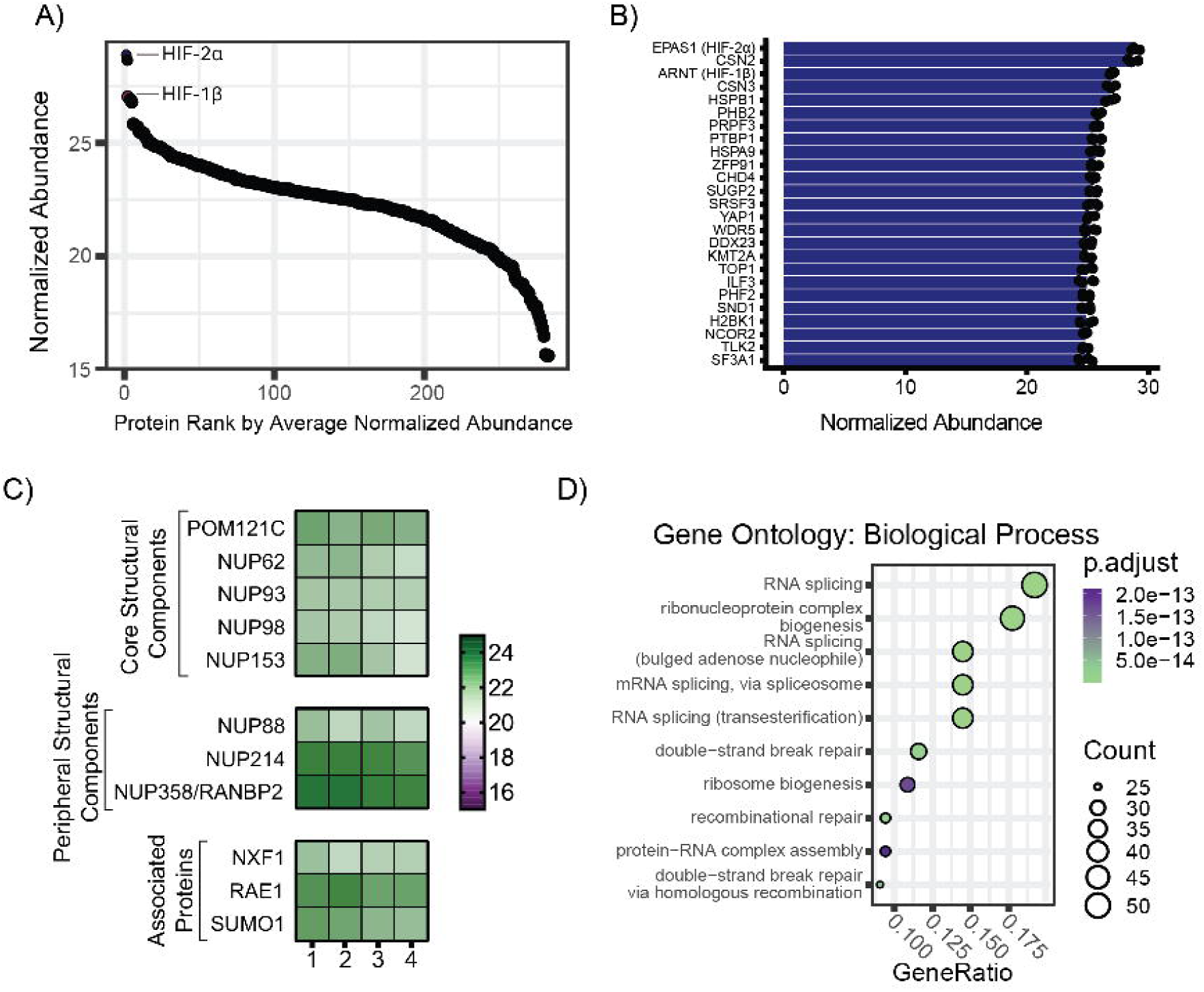
TurboID-MS expands the HIF-2α interaction network to include mRNA export factors. **A)** Rank plot displaying the abundance distribution of proteins identified in the HIF-2α TurboID-MS dataset. Proteins ranked by average normalized abundance. **B)** Top 30 ranked proteins identified by HIF-2α TurboID-MS are shown based on normalized abundance values. **C)** Heatmap depicting components of the Nuclear Pore Complex. **D)** Gene Ontology (GO) biological process enrichment analysis of TurboID-MS-identified HIF-2α-associated proteins. Dot size indicates the number of proteins contributing to each enriched term, while color represents adjusted p-value.

## Independent transcriptomic and genomic analyses support the biological relevance of the HIF-2α interactome

To further assess the biological relevance of our HIF-2α interactome, we integrated our proteomic dataset with publicly available proteomic and cancer genomic datasets. At the protein level, nearly all of the HIF-2α interactors identified here were detected in breast tumor proteomes from the Clinical Proteomic Tumor Analysis Consortium (CPTAC)^37^ (**Figure 5A**). Compared to a randomly identified set of 2,000 protein-coding genes which, on average, are identified in 50% of the tumor profiles, our HIF-2α interactome is identified in 89-98% of surveyed CPTAC tumor proteomes. Given the established role of HIF-2α in clear cell renal cell carcinoma (ccRCC),^3,13^ we also examined the presence of our interactors within a ccRCC proteome^38^. There, 96% of HIF-2α interactors were detected, compared with only 53% of randomly selected background proteins (**Figure 5B**). To test whether the HIF-2α interactors are differentially represented in tumor tissue, we compared each interactor against its own matched normal kidney in 232 ccRCC pairs profiled by quantitative mass spectrometry^39^. A large proportion of the interactors were identified in tumors more often than in normal tissue. In a typical patient, the share of HIF-2α interactors detected at the protein level is 3 percentage points (pp) higher in tumor than in matched normal (**Figure S4A**). We evaluated whether the expression level of these interactors differ when they are identified in both tumor and normal tissue from the same patient. Among interactors quantified in both tissues of at least 10 patients, paired abundance was shifted upward in tumor, significantly exceeding the background distribution (**Figure 4SB**), indicating that proteins recovered in proximity to HIF-2α are coordinately enriched in ccRCC tumor tissue.

**Figure 5:**
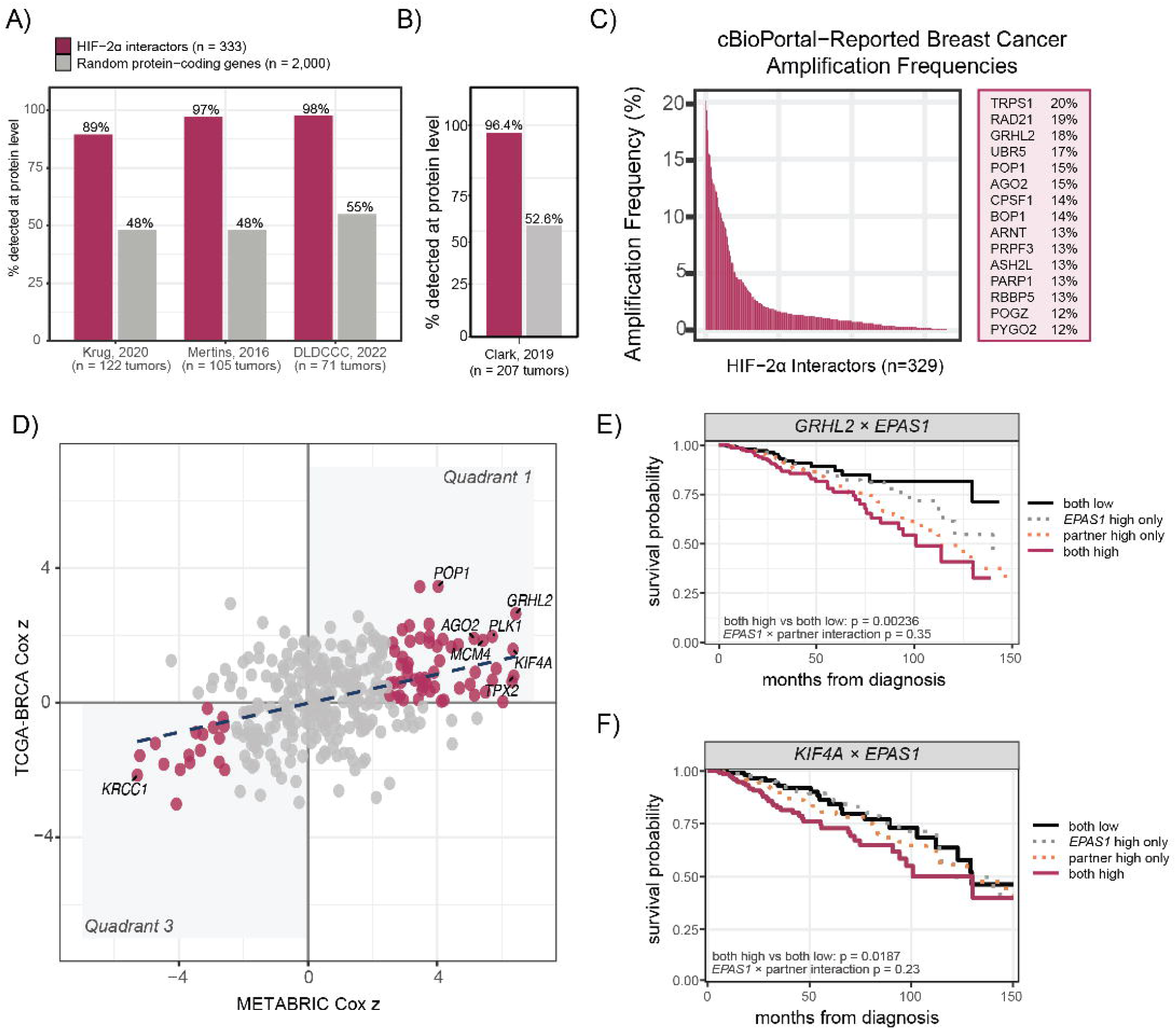
Independent transcriptomic and genomic analyses support the biological relevance of the HIF-2α interactome. **A)** The proportion of our HIF-2α interacting proteins that have been identified in three separate breast cancer tumor proteomics investigations. A control group of 2,000 protein-coding genes was randomly generated and the proportion of those proteins’ detection in the same datasets are plotted alongside our interactors. **B)** The proportion of our HIF-2α interacting proteins that have been identified in a ccRCC tumor proteome. **C)** Frequency of copy number amplification of HIF-2α proteomic interactors in breast cancer determined using cBioPortal Oncoprint. Bars represent the percentage of breast tumors harboring amplification of each indicated interactor that coincides with EPAS1 amplification. **D)** Comparison of the genes encoding our proteomic interactors’ ability to predict patient survival from two distinct breast cancer datasets: TCGA PanCancer and Metabric. **E)** Survival curve analysis of EPAS1, the gene encoding HIF-2α, and GRHL2 as well as their interaction. **F)** Survival curve analysis of EPAS1, the gene encoding HIF-2α, and KIF4A as well as their interaction.

We next asked whether the HIF-2α interactors detected in breast tumors are clinically relevant. We first examined copy-number amplification of the identified HIF-2α interactors using a breast cancer dataset from cBioPortal. Among our HIF-2α interactors, *TRPS1*, *RAD21*, *GRHL2*, and *UBR5* are frequently amplified in breast cancer (**Figure 5C**). We ran the same analysis for the ccRCC cancer dataset on cBioPortal and found a distinct series of commonly amplified proteins (**Figure S4C**). Although frequent amplification suggests that these genes may act as oncogenes, this analysis alone cannot show whether their interaction with HIF-2α contributes to that role. To ascertain more robust conclusions on these proteins’ mechanistic role in cancer, we took two independent gene expression datasets from breast cancer patients, TCGA Pan-Cancer Breast Cancer and Metabric, and compared the ability of our interactors to predict patient outcomes in breast cancer (**Figure 5D**). Interactors whose gene expression was consistently associated with survival in both independent datasets fall in quadrants one and three. Quadrant one contains interactors whose higher expression was associated with worse survival in both cohorts. Next, we evaluated the interaction effect of high expression of *EPAS1* (the gene encoding HIF-2α) as well as the top quadrant one interactors on patient survival probability. In both instances, high mRNA expression of both *EPAS1* and either *GRHL2* (**Figure 5E**) or *KIF4A* (**Figure 5F**) predicts worse patient outcomes than low expression of both. However, the interaction effect is largely driven by the predictive nature of the interactor (*GRHL2* or *KIF4A*) alone. These two proteins have distinct roles in cancer progression: GRHL2 acts as a transcriptional regulator of epithelial identity, competing with ZEB to restrain epithelial-mesenchymal transition^40,41^ and, in breast cancer, rewiring estrogen receptor-dependent enhancer activity, whereas KIF4A is a mitotic chromokinesin whose overexpression drives proliferation and invasion and tracks with advanced stage, higher grade, and shorter survival in ccRCC^42,43^. Although these analyses cannot show that HIF-2α’s interactions with these proteins drive breast cancer progression, they highlight candidates such as *GRHL2* and *KIF4A* for future functional studies. Collectively, these orthogonal analyses indicate that a subset of the identified HIF-2α interactors may be clinically relevant in breast and kidney cancers.

## Discussion

Protein-protein interactions are fundamental to cellular function, enabling proteins to assemble into functional complexes and integrate regulatory pathways. This is particularly important for transcription factors whose activity is often governed by interactions with other transcription factors, chromatin regulators, transcriptional coactivators and corepressors, and components of the transcriptional machinery. Defining these interaction networks can therefore provide insight into the mechanisms regulating transcription factor activity and function. Pulldown-based proteomics is a starting point to elucidate these functional interaction networks. In this study, we carefully evaluated HIF-2α’s baseline proteomic interactome in breast cancer through IP-MS and expanded these findings with TurboID-MS. In general, our two pulldown approaches enriched for proteins across similar biological functions, only differentiating in the exact protein members identified (**Figure S4D**). Our analyses uncovered new proximal interactors; TurboID-MS validated some of our findings in IP-MS as well as expanded our findings to suggest a role for HIF-2α in mRNA export through the identification of TREX components and nuclear pore complex proteins.

From this work the HIF-2α interactome has been expanded beyond canonical transcriptional regulators, revealing extensive associations with chromatin remodeling complexes, Mediator components, and epigenetic regulators that likely facilitate transcriptional activation. A common pathway identified across both pulldown methods was RNA splicing. Numerous spliceosomal proteins, including SF3A, SF3B, PRPF, PLRG1, CDC5L, RBMX2, RBM22, SON, and SRRM1, together with multiple RNA helicases (DDX1, DDX20, DDX24, DDX46, DDX47, DDX56, DHX16, and DDX23) and RNA-binding proteins (CELF1, MBNL1, PUM3, CIRBP, AGO2, and RBM family proteins), were identified across the combined datasets. Given recent work which suggests that HIF-2α helps drive DNA associations with nuclear speckles^17^, this finding is very compelling.

In conclusion, this work suggests that IP-MS and TurboID-MS are powerful when used in conjunction to gain a more comprehensive interactome. Collectively, these findings support a model in which HIF-2α functions as a central organizer of nuclear gene expression, coupling chromatin regulation, transcription, RNA processing, and mRNA export to optimize the cellular response to hypoxia. Publicly available clinical datasets provide supporting evidence that our interactors were more likely to be found within tumors when compared to a background set of randomly sampled proteins. Furthermore, for interactors that were identified across both tumor and normal tissue of the same patient, the interactors as a group were often more highly expressed in the tumor proteome than in the normal proteome. Further, several of these interactors are associated with patient survival in breast cancer, yet neither GRHL2 nor KIF4A has been investigated as a HIF-2α co-regulator.

This integrated resource provides a framework for future mechanistic studies aimed at defining how individual HIF-2α interactions contribute to normal hypoxic adaptation and to HIF-2α-driven diseases such as clear cell renal cell carcinoma. While we believe this work is an abundant resource for groups studying the biochemistry of HIF-2α, we acknowledge that one major limitation of our work is the lack of hypoxia treatment to our MCF7 cell lines. Here, HIF-2α was overexpressed with a double proline mutation which stabilizes it against VHL degradation which permitted a baseline characterization of the HIF-2α interactome. Future work would benefit from the additional variable of hypoxic conditions to capture the interaction landscape of endogenous wildtype HIF-2α.

## Acknowledgements

The authors would like to thank Abhisha Sawant Dessai for her input on this project.

## Author Contributions

C.L.F., K.A.A., and S.L.B. conceived the project with input from K.A.T. and C.R.G.

C.L.F., K.A.A., S.L.B., and C.R.G. designed the experiments.

C.L.F., A.K., and F.N.D.L.V. performed experiments and collected the data.

C.L.F. analyzed the data with help from H.H., C.R.G., M.B.G., and B.A.G.

C.L.F. prepared figures with input from C.R.G., R.Y., and S.L.B.

C.L.F. wrote the manuscript, with input and editing from C.R.G. and S.L.B.

All authors reviewed the manuscript.

---

**Supplemental Figure 1:**
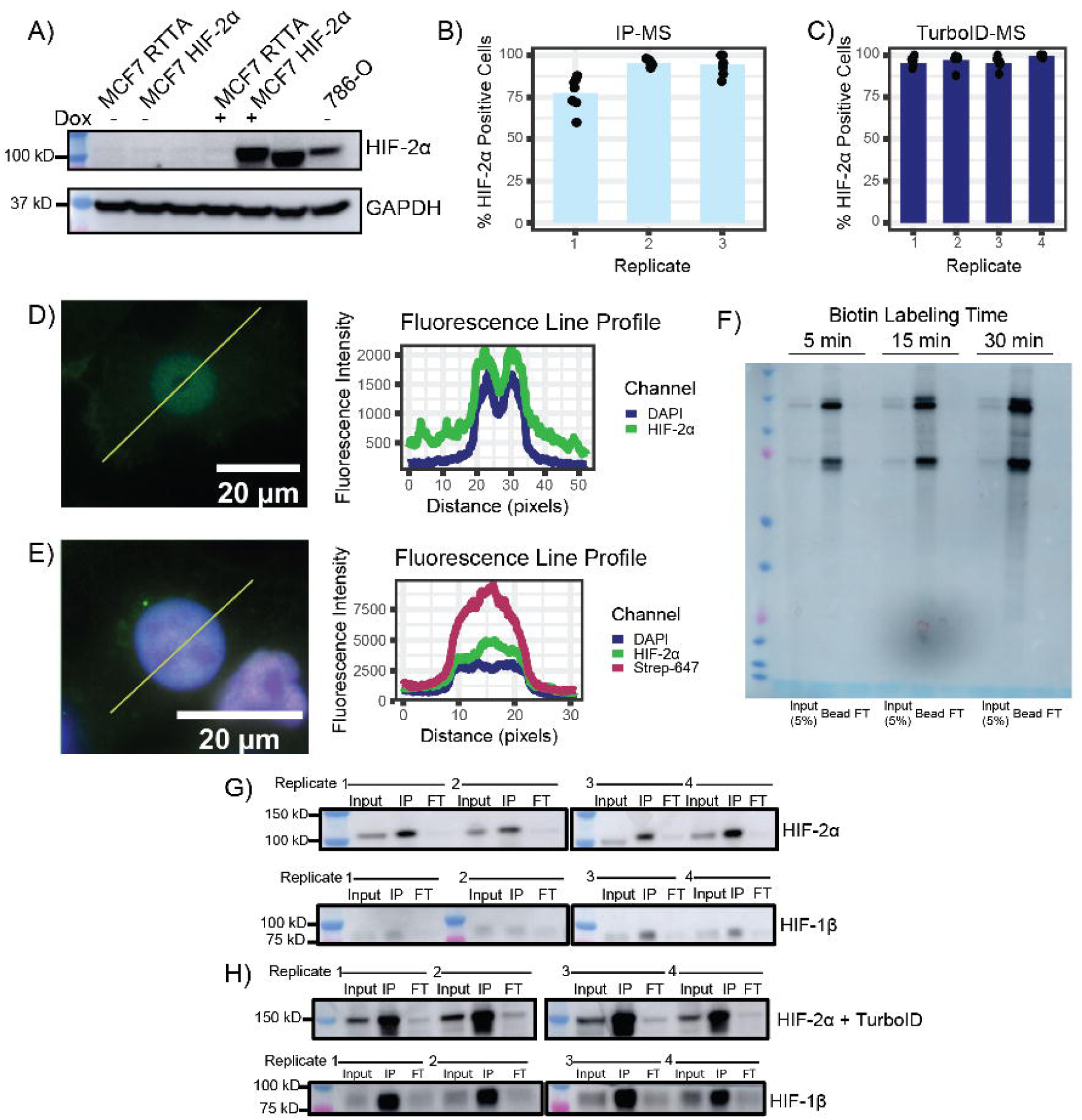
**A)** Western Blot image of MCF7-HIF-2α cell lysates with and without doxycycline treatment. Lanes include MCF7-rtTA (parental line lacking the HIF-2α lentiviral construct, MCF7-HIF-2α with and without doxycycline, and 786-O cells, which express HIF-2α endogenously. Only relevant lanes labeled. **B)** Quantification of HIF-2α positive staining of the MCF7-HIF-2α cell lines shown in Figure 1D. **C)** Quantification of HIF-2α positive staining of the MCF7-HIF-2α-TURBO cell lines shown in Figure 1E. **D**) Fluorescence intensity along a line drawn through one representative MCF7-HIF-2α cell, plotted for each channel acquired. **E)** Fluorescence intensity along a line drawn through one representative MCF7-HIF-2α-TurboID cell, plotted for each channel acquired. **F)** Western blot of MCF7-HIF-2α-TurboID cells labeled with biotin for 5, 15, or 30 minutes. Biotinylated protein was detected with streptavidin-HRP in the streptavidin-bead-captured ("Bead") fraction at each timepoint. **G)** Western blot of the same cells used for the IP-MS proteomics samples, probed for HIF-2α and HIF-1β. Three fractions per sample were collected: input, immunoprecipitate (IP), and flow-through (FT), loaded at 2.5%, 25%, and 2.5% of total, respectively. **H)** Western blot of the same cells used for the TurboID-MS proteomics samples, probed for HIF-2α and HIF1β,with fractions loaded at the same percentages as in G).

**Supplemental Figure 2:**
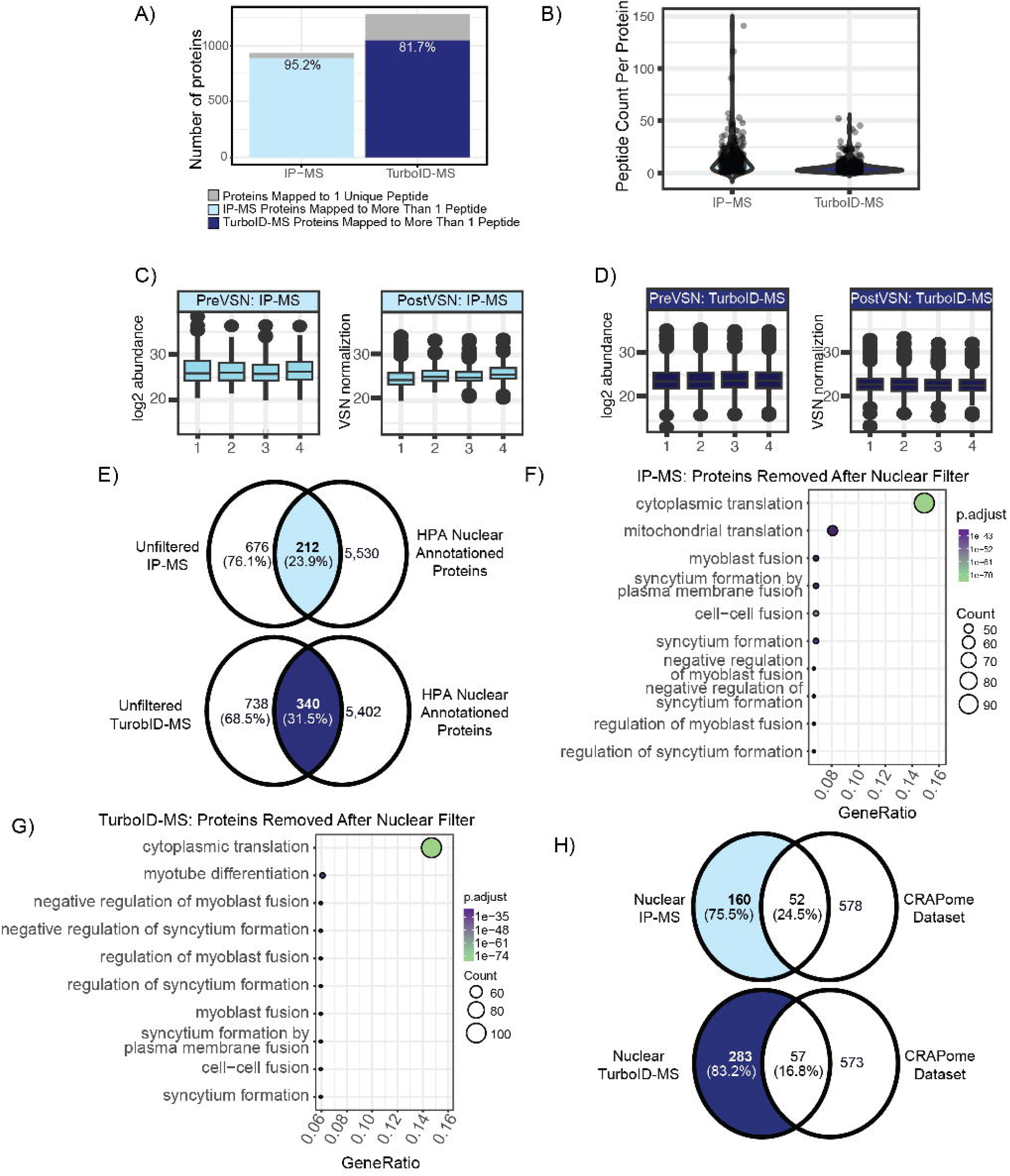
**A)** Stacked Bar Plot representation of fraction of proteins mapped to 1 unique peptide or more than 1 unique peptide (light blue denotes IP-MS and dark blue denotes TurboID-MS datasets). **B)** Violin plots of the peptide counts for all proteins with more than one unique peptide. **C)** Pre and post VSN normalization abundance spreads for IP-MS data. **D)** Pre and post VSN normalization abundance spreads for TurboID-MS data. **E)** Venn diagram representation of dataset filtering for both datasets: proteins that were not annotated by the Human Protein Atlas (HPA) were removed. **F)** The Gene Ontology - Biological Process categories for the 676 proteins filtered out of the IP-MS dataset due to their absence of nuclear annotation by the HPA. **G)** The Gene Ontology - Biological Process categories for the 738 proteins filtered out of the TurboID-MS dataset due to their absence of nuclear annotation by the HPA. **H)** Venn diagram representation of filtering of proteins found in CRAPome dataset CC156 in both datasets.

**Supplemental Figure 3:**
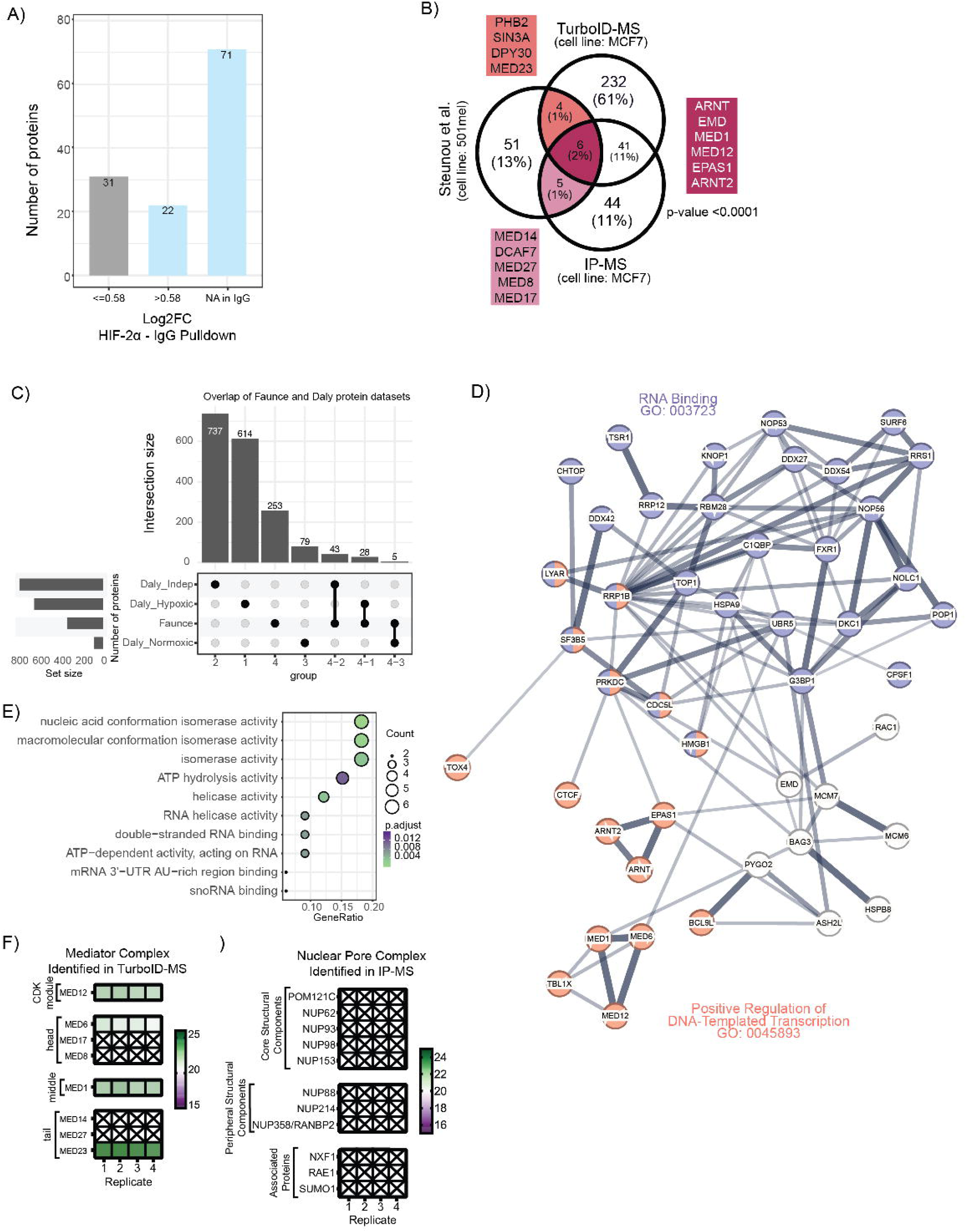
**A)** Bar plots representing fold change comparisons between HIF-2α pulldown abundance and IgG pulldown abundance. Proteins that had a log2FC <= 0.58 (IgG high) are represented in grey. Proteins that either were not detected in IgG or were >0.58 enriched over IgG are represented in light blue. **B)** Venn diagram of the overlap between our datasets and the previously published HIF-2α interactome by Steunou et al. Statistical significance of the overlap was determined using a hypergeometric test. **C)** UpSet plot representation of the intersections between our combined dataset (IP-MS and TurboID-Ms proteins) and Daly et al. Daly et al.’s published datasets are segmented as oxygen independent (Daly_Indep), 1% oxygen dependent (Daly_Hypoxic), and 21% oxygen dependent (Daly_Normoxic). **D)** StringDB analysis of the 74 commonly identified proteins across both methods. Notable pathways include RNA binding (p-value = 1.89e-15) and positive regulation of DNA-templated transcription (p-value = 5.83e-5). **E)** Gene Ontology – Molecular Function analysis of the commonly identified proteins between IP-MS and TurboID-MS. **F)** A heatmap of all mediator complex subunits found across both pulldown methods filled with abundance values from TurboID-MS. **G)** A heatmap of all nuclear pore complex subunits and associated proteins found across both pulldown methods filled with abundance values from IP-MS.

**Supplemental Figure 4:**
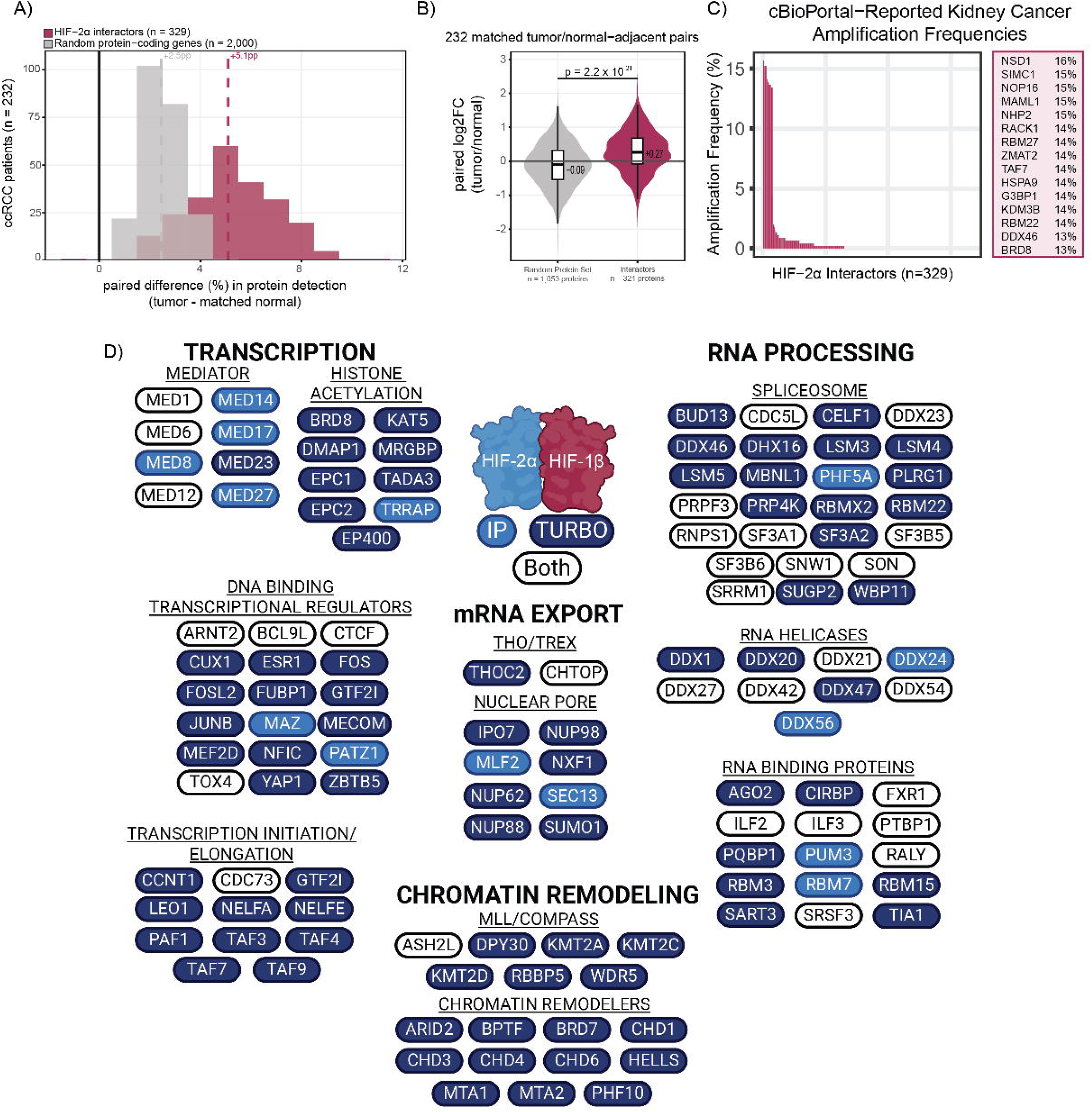
**A)** For each of the matched tumor / normal-adjacent pairs in the ccRCC proteomics data, the percentage of genes encoding proteins (either from our dataset or from a randomly selected set of 2,000 protein coding genes) quantified in the tumor sample minus the percentage quantified in the matched normal sample (percentage points, pp). Each patient contributes one value per set; dashed lines mark set medians (+2.5 pp for random genes, +5.1 pp for interactors). Positive values indicate that more proteins of that set were detected in the tumor than in its matched normal. The comparison is over presence/absence of a quantified value and is insensitive to how abundant a detected protein is. **B)** For each protein (either interactor from our dataset or a protein from the random set of protein-encoding genes) quantified in at least 10 pairs, the median across patients of the paired log2 ratio (tumor / normal-adjacent). Each point in the distribution is one protein, not one patient; medians annotated: −0.09 random, +0.27 interactors) Groups were compared by a two-sided Wilcoxon rank-sum test (p = 2.2 × 10⁻^21^; n = 715 random and 246 interactor proteins). Proteins missing in either member of a pair contribute no value, so this panel describes only the quantifiable subset. **C)** Frequency of copy number amplification of HIF-2α proteomic interactors in breast cancer determined using cBioPortal Oncoprint. Bars represent the percentage of ccRCC tumors harboring amplification of each indicated interactor that coincides with EPAS1 amplification. **D)** Cartoon summary of notable molecular function groups identified across both methods. White denotes identification across both approaches, light blue denotes identification in IP-MS only, and navy blue denotes identification in TurboID-MS only.

Supplemental Table S1: Table of normalized protein abundances for all biological replicates across both detection methods.

